# Easy, fast and reproducible Stochastic Cellular Automata with chouca

**DOI:** 10.1101/2023.11.08.566206

**Authors:** Alexandre Génin, Guillaume Dupont, Daniel Valencia, Mauro Zucconi, M. Isidora Ávila-Thieme, Sergio A. Navarrete, Evie A. Wieters

## Abstract

Stochastic cellular automata (SCA) are models that describe spatial dynamics using a grid of cells that switch between discrete states over time. They are widely used to understand how small-scale processes scale up to affect ecological dynamics at larger spatial scales, and have been applied to a wide diversity of theoretical and applied problems in all systems, such as arid ecosystems, coral reefs, forests, bacteria, or urban growth.

Despite their wide applications, SCA implementations are often ad-hoc, lacking performance, guarantees of correctness and poorly reproducible. *De novo* implementation of SCA for each specific system and application also represents a major barrier for many practitioners. To provide a unifying, well-tested technical basis to this class of models and facilitate their implementation, we built chouca, an R package that translates definitions of SCA models into compiled code, and runs simulations in an efficient way.

chouca supports SCA based on rectangular grids where transition probabilities are defined for each cell, with performance typically two to three orders of magnitude above typical implementations in interpreted languages (e.g. R, Python), all while maintaining an intuitive interface in the R environment. Exact and mean-field simulations can be run, and both numerical and graphical results can be easily exported.

Besides providing better reproducibility and accessibility, a fast engine for SCA unlocks novel, computationally intensive statistical approaches, such as simulation-based inference of ecological interactions from field data, which represents by itself an important avenue for research. By providing an easy and efficient entry point to SCAs, chouca lowers the bar to the use of this class of models for ecologists, managers and general practitioners, providing a leveled-off reproducible platform while opening novel methodological approaches.

## Introduction

The analysis of spatial patterns has proven essential to understand ecological system dynamics, and various modelling approaches have helped ground empirical patterns into ecological theory. Among such approaches, models based on Stochastic Cellular Automata (hereafter SCA), also called Probabilistic or Random Cellular Automata, or Locally-interacting Markov Chains, have been a particularly useful, heuristic and widely used approach (Wolfram, 1984; Louis & Nardi, 2018). Cellular automata are based on a grid of cells that switch over time between a finite number of states. Most often, SCA are considered over a rectangular grid, though other geometries can exist (van Baalen, M., 2000). A famous deterministic cellular automaton (CA) is Conway’s game of life, which is defined by two discrete states (“dead” and “alive”) and a set of deterministic rules to make cells switch between them (Gardner, 1970; Bays, 2010). Stochastic cellular automata follow the same principles, but state transitions occur with a given probability instead of being based on deterministic rules. The probabilities of a cell switching from one state to another is assumed to depend on model parameters, the global state of the system (the proportion of cells in each state), and the local neighborhood of the focal cell. In all cases, the system future dynamics is probabilistically defined by its current state, *i*.*e*. dynamics are memoryless.

The use of SCA in ecology is widespread, as they have been used to describe the dynamics of a large array of ecosystems, including mussel beds (Guichard et al., 2003), arid ecosystems (Kéfi et al., 2007), forests (Heinonen & Pukkala, 2007), rocky shores (Wootton, 2001), coral reefs (Muthukrishnan et al., 2016; Génin et al., 2024) and plant communities (Lanzer & Pillar, 2002). SCA are often used to understand how local processes can scale up to affect landscape-wide properties, such as the persistence or extinction of a given species, the type of spatial patterns arising at the scale of a landscape (Pascual et al., 2002), or the spread of fire or epidemics. The latter case is a classical application of SCA in applied ecology, where data on local processes affecting forest stands can be coupled with GIS data to provide guidance on forest fire sensitivity (Yassemi et al., 2008). Though common in ecosystem modelling, and the primary purpose for this package, SCA are useful beyond this sole discipline, and are for example also used to describe epidemics (Yakowitz et al., 1990; Keeling, 2000), tumor growth (Moreira & Deutsch, 2002) or organism development (Manukyan et al., 2017).

The popularity of SCA is probably rooted in their relatively light need for formal mathematics compared to other approaches modelling spatial processes (*e*.*g*. partial derivative equations): only the probabilities of transitions between states need to be defined. The drawback of this simplicity is that the numerical simulation of SCA is typically computationally-intensive. Current approaches to do so efficiently rely on approximations, either assuming no spatial structure (mean field approximation), or using approximations to obtain analytical results, such as spatial moment equations (Lion, 2016; of which pair approximation is a specific case; Matsuda et al., 1992; Iwasa, 2000). However, these approximations are inappropriate when long-range correlations occur within a landscape (Iwasa, 2000), or when the full simulated landscape is needed as a model output, for example to compute spatial metrics (Génin et al., 2018). In many cases, the explicit numerical simulation must be run, which is often done on grids that may not be large enough to fully capture emergent spatial patterns (van de Koppel et al., 2011; Majumder et al., 2021). On top of these performance issues, most ecological studies based on SCA come with their own implementation. This opens the possibility of errors in code and often makes it difficult to reproduce model simulations. We aim at alleviating these issues with `chouca`, which provides a unifying and well-tested technical basis to SCA. Our goal is to improve the performance and accessibility of this class of models, and ultimately allow ecologists to spend more time exploring the behavior of their models, rather than on their implementation.

### Supported models

The R package `chouca` works with 2-dimensional rectangular grids of cells (a “landscape”). Each cell can be in one of a finite set *S* of *n* elementary states *S*_1_ … *S*_*n*_. Probabilities of transition are assumed to depend only on (i) the proportion of neighbors in each state, captured by the vector ***q*** = (*q*_1_, …, *q*_*n*_), (ii) the proportion of cells in a given state in the whole landscape, ***p*** = (*p*_1_, …, *p*_*n*_), and (iii) a set of constant model parameters ***θ***.

`chouca` has been primarily designed for modeling the dynamics of sessile organisms over space, which do not move and reproduce through the dispersal of propagules (*e*.*g*. gap-models; Shugart & West, 1981). This includes a wide range of organisms, ranging from forests, herbaceous plants or coral reefs, but the supported models may also be useful to describe other ecological situations. For the moment, the types of models that can be implemented exclude cellular automata in which an intermediate distance of interaction is considered (*e*.*g*. through a dispersion kernel; Muthukrishnan et al., 2016), or those in which a preferential direction exists (*e*.*g*. modeling water redistribution on a slope; Mayor et al., 2013), though these limitations are planned to be lifted in later versions. More importantly, SCA that are based on the dynamics of pairs of cells, for instance in which two cells swap their respective state, cannot be implemented directly. This is typical when representing the movements of an organism in a landscape (Pascual et al., 2002), and we provide links to alternative software at the end of the article for these models.

In the package, the probabilities of transition of a cell from state *i* to *j, P*(*S*_*i*_ → *S*_*j*_), can be any function of ***p, q***, and ***θ*** – however, it will work best and without approximation if it has the following form:

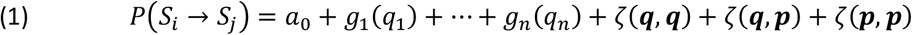

where, for any transition *S*_*i*_ → *S*_*j*_, *a*_0_ is a constant, *g*_*s*_ is any univariate function of *q*_*s*_, and *ζ*(***x, y***) is the sum, defined for two vectors ***x*** = (*x*_1_, …, *x*_*n*_) and ***y*** = (*y*_1_, …, *y*_*n*_) as

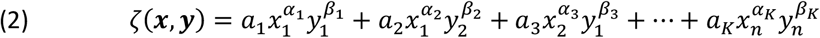

in which the (*a*_*i*_)_*i*∈[1:*K*]_, (*α*_*i*_)_*i*∈[1:*K*]_ and (*β*_*i*_)_*i*∈[1:*K*]_ are constants and *K* is the total number of terms of the sum. In practice, this functional form is flexible enough to approximate the probabilities of transition of many ecological models.

Implementing and working with an SCA in `chouca` typically consists in four steps, in which the user (1) defines the model states and the transitions between them, (2) creates an initial landscape (grid of cells), (3) runs the model, and (4) displays or extracts the results (Figure 1). We detail in this paper this workflow (Figure 1) – documentation is available throughout the package, individually for each function, or as a whole in an R “vignette”, accessible with the command `vignette(“chouca-package”)`.

**Figure 1.**
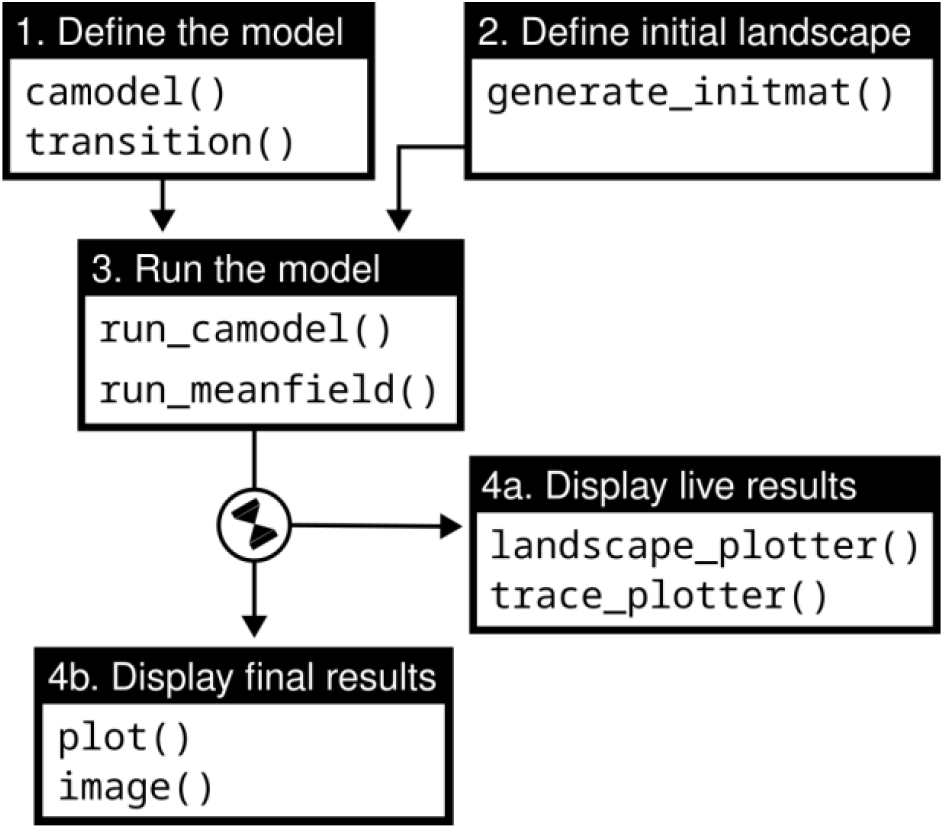
Main tasks (boxes) of the chouca package, and their associated sets of functions

### Example applications

#### A simple model of mussel bed

To illustrate how a stochastic cellular automaton can be defined with `chouca`, we use the model of Guichard et al. (2003), which describes the dynamics of mussels colonizing rocks exposed to waves. This model has three cell states (i) “disturbed”, (ii) “empty” and (iii) “occupied” (by mussels). During a single time-step, a disturbed cell becomes an empty cell with probability 1. Such transition can be defined by using a call to the R function `transition()`:

~~~
transition(from = “disturbed”, to = “empty”, ∼ 1)
~~~

This statement declares a transition from a “disturbed” state to an “empty” state, with the last argument being a symbolic expression starting with “`∼`”, that describes how to compute the probability, here being simply equal to the constant “1”.

The model assumes that the establishment of new individuals always occurs next to existing mussels. In other words, for a focal cell in the “bare” state, its probability of switching to the “mussel” state is not constant, but given by *αq*_*mussel*_, where *α* is a productivity rate, and *q*_*mussel*_ is the proportion of cell neighbors in the mussel state. Such transition is defined by the following call in R:

~~~
transition(from = “empty”, to = “mussel”, ∼ alpha * q[“mussel”])
~~~

Here, `q[“mussel”]` is used to refer to the proportion of cell neighbors in the “mussel” state, a continuous number between 0 and 1. Similarly, `p[“mussel”]` can be used to refer to the global proportion of cells in the landscape in the “mussel” state.

Mussels can be perturbed by incoming waves, which dislodge them and turn them into “disturbed” cells. In this model, the probability of a mussel cell to become disturbed is the sum of a baseline term *δ*, and an additional term *d*, which is non-zero only if the mussel cell has one or more disturbed neighbors:

~~~
transition(from = “mussel”, to = “disturbed”, ∼ delta + d * (q[“disturbed”] > 0))
~~~

The original model considers that cells are neighbors when they share an edge (current options include a 4-way or von-Neumann neighborhood, the other option being a Moore or 8-way neighborhood), and uses a toric space for simulations, meaning that the up/leftmost cells of the grid are neighbors of the bottom/rightmost cells. Putting everything together, this model can be defined using the following syntax:

~~~
musselbed_mod <-camodel(
  transition(from = “disturbed”, to = “empty”, ∼ 1),
  transition(from = “empty”, to = “mussel”, ∼ alpha * q[“mussel”]),
  transition(from = “mussel”, to = “disturbed”, ∼ delta + d * (q[“disturbed”] > 0)),
  parms = list(alpha = 1, delta = 0.2, d = 1), # parameters
  wrap = TRUE, # toric space
  neighbors = 4 # 4-way neighborhood
)
~~~

The call to the `camodel()` function above estimates the constants in the functional form described in equation 1 to match the parameters of the model. If this process yields large residual error, for example because the transition probabilities do not correspond to the functional form in equation 1, a warning is produced. The structure of the model can be displayed on the R console using `print()`, or as a graph using the generic function `plot()` (Figure 2).

**Figure 2.**
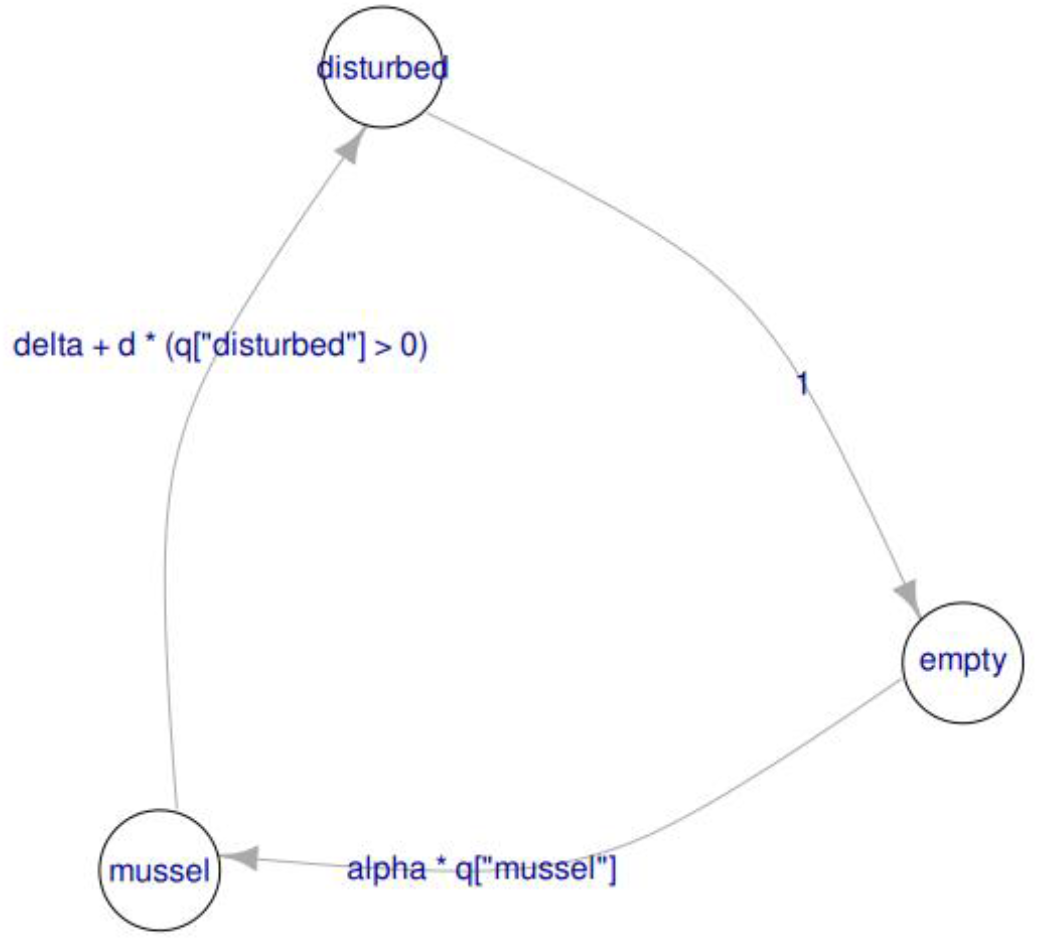
Structure of the mussel bed model displayed as a graph using plot(musselbed_mod), in which nodes are states, and arrows represent the possible transitions, with the expression used to compute their probabilities

~~~
> print(musselbed_mod)
Stochastic Cellular Automaton
States: disturbed empty mussel
Transition: disturbed -> empty
  ∼ 1
Transition: empty -> mussel
  ∼ alpha * q[“mussel”]
Transition: mussel -> disturbed
  ∼ delta + d * (q[“disturbed”] > 0)
Neighborhood: 4×4
Wrap: TRUE
Max error: 0 (OK)
Max rel error: 0 (OK)
~~~

Once the model is created, an initial landscape can be defined with a given number of rows and columns using `generate_initmat()`, which creates a landscape with randomly-distributed cell states in space, but following the specified initial covers:

~~~
init_landscape <-generate_initmat(musselbed_mod,
                                  pvec = c(disturbed = 0.1, empty = 0.1, mussel = 0.8),
                                  nrow = 64, ncol = 64)
~~~

The model can then be simulated for a given number of time steps, below from 0 to 50, using `run_camodel()`. Standard methods such as `plot p()` or `image()` can be used to display the global covers after the model has run, or the resulting landscapes:

~~~
output <-run_camodel(musselbed_mod, init_landscape, times = seq(0, 50))
plot(output)
image(output)
~~~

By default, `chouca` uses a C++ backend based on Rcpp (Eddelbuettel, 2013), which has reasonable performance. This can be improved by compiling the model code just before the simulation is run, and using memoization so that transition probabilities are computed only once for cells with the same neighborhood configuration. This typically improve performance by two orders of magnitude over a typical implementation (Figure 3; Schneider et al., 2016), which can be further increased by parallelizing computations over multiple cores, though this is a less efficient approach. Enabling these options can be done by passing control arguments as an R list object:

**Figure 3.**
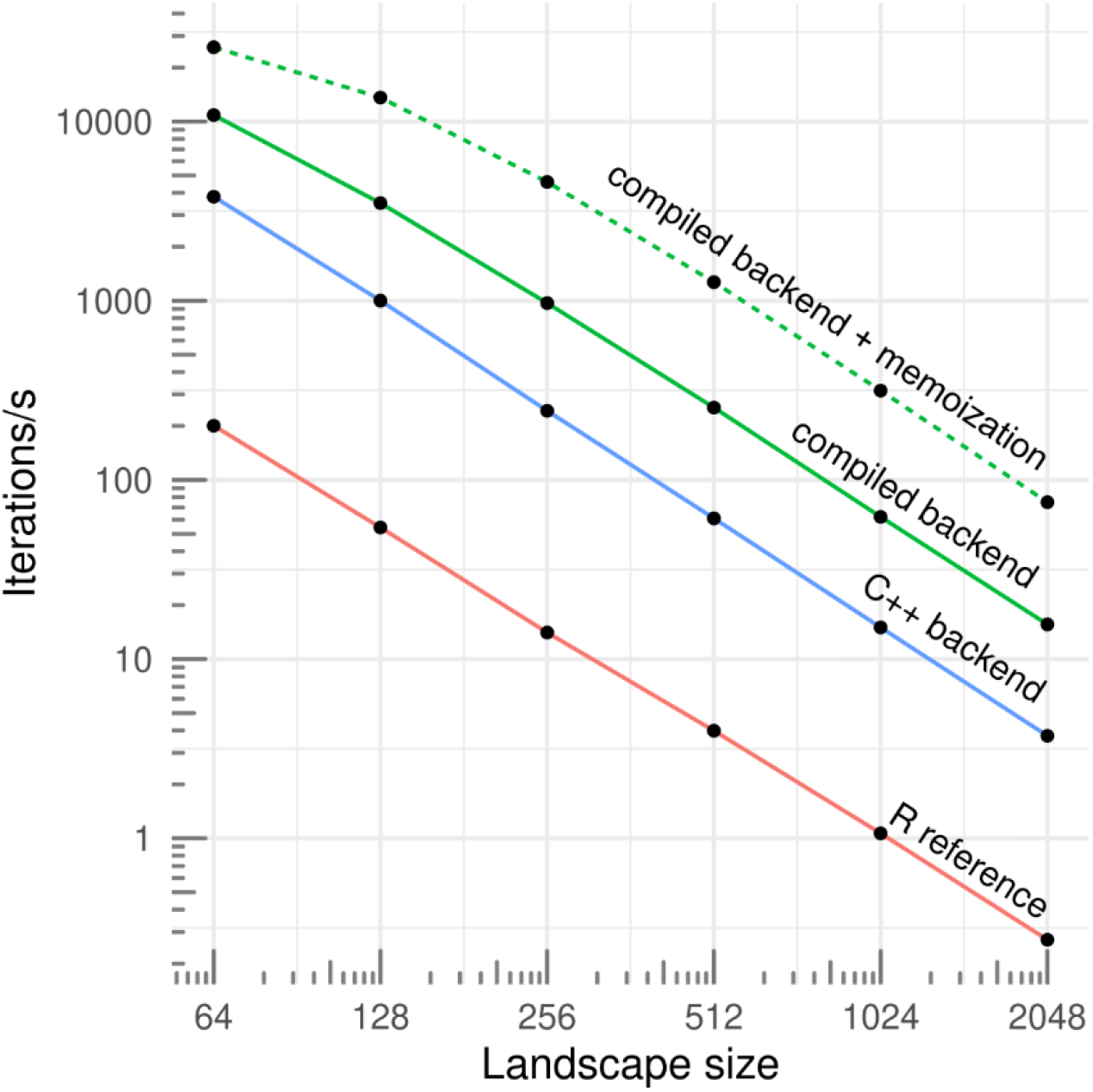
Simulation speeds for a set of three simple ecological models (2-3 states and 3-4 transitions) according the the grid size, for a pure-R implementation (R reference; Schneider et al., 2016), and three backends provided by chouca (blue and green lines). Single-core performance on a 2020 desktop computer.

~~~
control_args <-list(engine = “compiled”,
                    cores = 4,
                    precompute_probas = TRUE)
output <-run_camodel(musselbed_mod, init_landscape, niter = 256, control = control_args)
~~~

This “control” list defines how the simulation is run, which data to save from the simulation, or the textual output to print while the simulation is running (a complete list of options can be found in the help page by using `? run_camodel` on the R console). The user can also supply custom functions that will be run as the simulation is running. This can be useful, for example, to display landscapes, covers, or compute statistics on the 2D landscape as the simulation is running, which we illustrate in the following sections.

#### Graphical explorations

Because an SCA describes dynamics over landscapes, they are particularly well-adapted to pattern-oriented modelling (Grimm et al., 1996), in which a model is defined and revised based on a qualitative or quantitative comparison with empirical patterns. Likelihood-based approaches are increasingly popular to compare models with data (Hartig et al., 2011 and section below), but the qualitative comparison and visual exploration of model dynamics remains an essential phase for spatial models. To make this modeling step more accessible, we made it easy to investigate visually the behavior of models, and illustrate here this approach with an epidemiological example.

Keeling (2000) uses an SCA-based approach to investigate the spread of a parasite over space, with an application to forests. This very simple model uses three states, “host”, “parasitized”, “empty”, and can be defined as follows in `chouca:`

~~~
mod <-camodel(transition(from = “empty”, to = “host”,
                         ∼ 1 - (1 - g)^(4 * q[“host”])),
              transition(from = “host”, to = “parasitized”,
                         ∼ 1 - (1 – T)^(4 * q[“parasitized”])),
              transition(from = “parasitized”, to = “empty”,
                         ∼ 1),
              parms = list(g = 0.05, T = 0.5),
              wrap = TRUE,
              neighbors = 4)
~~~

where *g* is the growth rate of the host, and *T* the transmissibility of the parasite (this model assumes that infection is always fatal, so the transition from “parasitized” to “empty” is equal to 1).

This model can produce interesting “epidemiological fronts”, which stem from the way the parasite spreads to its neighboring hosts, killing them and leaving behind empty, bare patches that are later recolonized by the host, albeit at a slower rate. This phenomenon of fronts propagating through an excitable media may be difficult to quantify formally, but can be easily visualized from model outputs. This can be done in `chouca` by setting the `run_camodel()` function to display the results as the simulation is run. To do so, we adjust the control list to include a function that displays the landscapes:

~~~
options <- list(custom_output_fun = landscape_plotter(mod),
                custom_output_every = 1)
~~~

then run the model on a 256×256 grid seeded with 10% of cells in the “parasitized” state:

~~~
initmm <-generate_initmat(mod, c(host = 0.9, parasitized = 0.1, empty = 0),
                          nrow = 256, ncol = 256)
output <-run_camodel(mod, initmm, times = seq(0, 1024), control = options)
~~~

The above lines of code will run the model and display the changing landscape, which allows investigating the spreading patterns of the parasite. Once this visualization step is done (Figure 4, and animated version in SM1), the options set to visualize the landscape can be removed to reduce simulation times, for example to investigate the model behavior along a range of parameter values.

**Figure 4.**
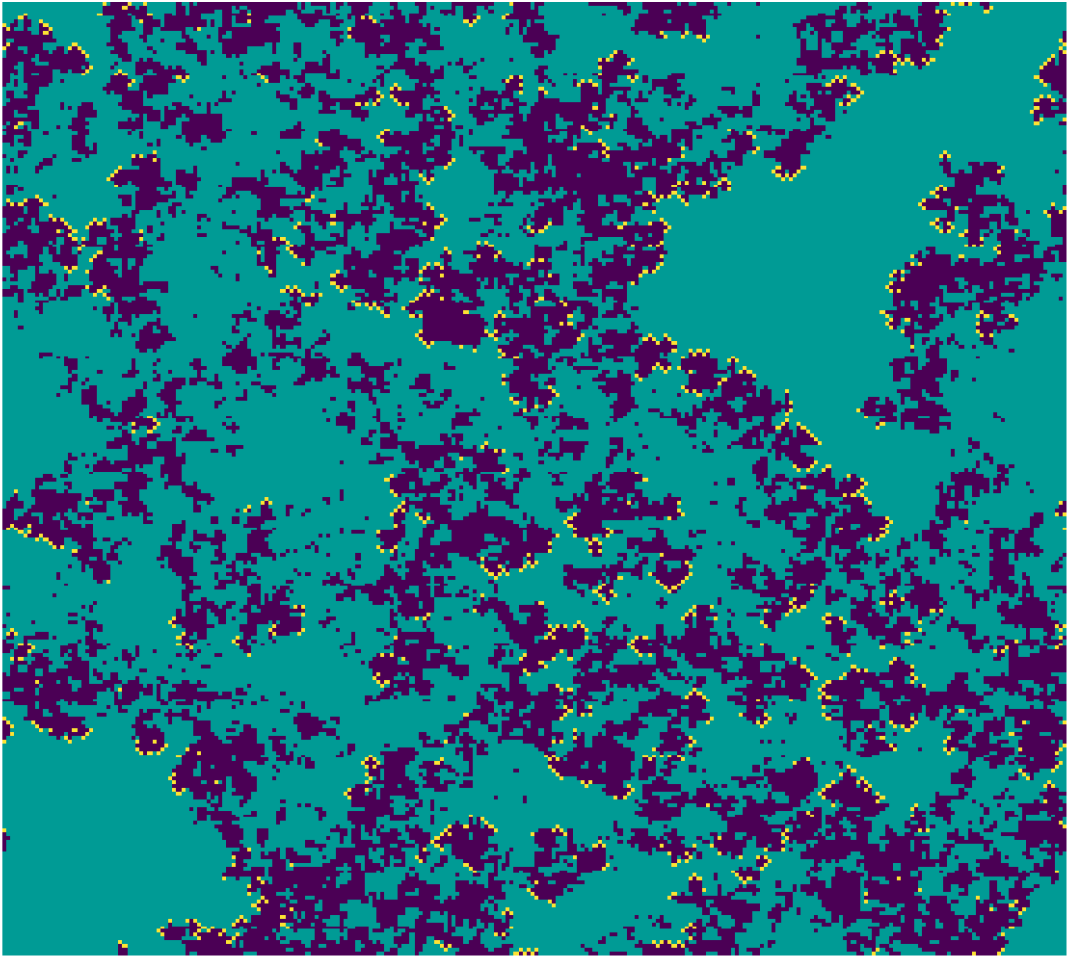
Landscape patterns of the host-parasitized-empty model (respectively green, yellow and blue) as displayed on screen.

#### Inference of local interactions from landscape-scale patterns

Because SCA are defined on grids, a natural application is to compare their output to empirical raster data, such as remote-sensing images, to infer local-scale ecological interactions from landscape-wide spatial patterns. Arid systems provide a good illustration of this approach: in those systems, interactions between plants are often a balance of negative effects, through competition for nutrients or water, with positive effects, for example with an increased survival of seedlings below the canopy of taller plants (Valiente-Banuet & Ezcurra, 1991). These effects can be strong enough for new plants to mostly establish below existing plants, which results in their aggregation into patches, and has important consequences for the resilience of those systems to changes in aridity (Kéfi et al., 2007). The sizes and numbers of those patches can be readily quantified from remote-sensing images, and such patterns can be used to infer whether facilitation occurs between plants (Xu et al., 2015; Chen et al., 2022). This is traditionally done by summarizing the spatial structure into spatial statistics, such as spatial autocorrelation (Sankaran et al., 2017) or type of patch size distribution (Siteur et al., 2023; Pichon et al., 2024) and linking the observed changes in those metrics to theoretical results (Scanlon et al., 2007; Kéfi et al., 2024). However, such qualitative comparison logically results in a qualitative and corroborative result, *i*.*e*. is there or is there not facilitation between plants, rather than a more informative quantitative result, *i*.*e*. how strong facilitation is between plants. A quantitative inference must be based on an approach that links quantitatively some aspects of the spatial patterns to the strength of facilitation. This can be done by defining a model that links the strength of facilitation with the expected patterns, and finding model parameters that maximize agreement between model output and data. This is expensive computationally, but this limitation is alleviated by a fast SCA engine such as chouca, as we show below.

We define here a model of an arid ecosystem with two states, “bare” and “vegetated”. A bare cell can become a vegetated cell with the probability *p*_*plant*_ . A vegetated cell can become bare (plants die) with the probability *d*(1 − *βq*_*plant*_), where *d* is a constant mortality rate, that is reduced by the coefficient *β. β* captures the local effect of plants, either facilitation when they increase the survival of plants near existing vegetation (*β* > 0), or competition when survival is decreased (*β* < 0). This model is implemented in `chouca` using:

~~~
facilitation_mod <-camodel(
  transition(from = “bare”, to = “plant”, ∼ p[“plant”]),
  transition(from = “plant”, to = “bare”, ∼ d * (1 – beta * q[“plant”])),
  parms = list(beta = 0.5, d = 0.85),
  wrap = TRUE,
  neighbors = 4
)
~~~

We ran the model till equilibrium on a 1024×1024 grid, to simulate a landscape that would be obtained from empirical data (*e*.*g*. a remote sensing image) using (*d, β*) = (0.85, 0.5). From this landscape used as observed data, we computed the distribution of pairs, which summarizes all the possible pairs of neighboring cells present in the landscape ***n***_***obs***_ = (*n*_*p*,0_, *n*_*p*,*p*_, *n*_0,0_). We then define the likelihood *P*(***n***_***obs***_│*d, β*) assuming these observed number of pairs follow a multinomial distribution of size *N*_*p*_ (the sum of all pairs, a fixed number given the grid size and neighborhood type), and probabilities ***μ*** = (*μ*_*p*,0_, *μ*_*p*,*p*_, *μ*_0,0_). ***μ*** defines the relative probabilities of observing each type of pair in the grid, which depend on the particular values of *d* and *β*, and can be estimated by simulating the landscape with these parameter values. This estimate of the likelihood formally quantifies the agreement between model and data, and allows exploring the parameter space in terms of *d* and *β* to find the most likely parameter combinations given the observed patterns.

We find that this approach can recover the parameter values used for *d* and *β*, with only one global maximum in the likelihood function (Figure 5; code provided in SM2), showing that the distribution of pairs is a sufficient statistic to recover the model parameters. Applying such an approach to empirical data would require further testing of model assumptions, for example, to investigate whether facilitation occurs on the recruitment of new plants instead of on the mortality of adult plants – this can be done by simply changing the model definition above. Because this approach is likelihood-based, model support can be compared using traditional statistics, such as AIC, and a Bayesian approach can be used to estimate credible intervals on parameter values, or use informative priors grounded in knowledge about the system (Hartig et al., 2011).

**Figure 5.**
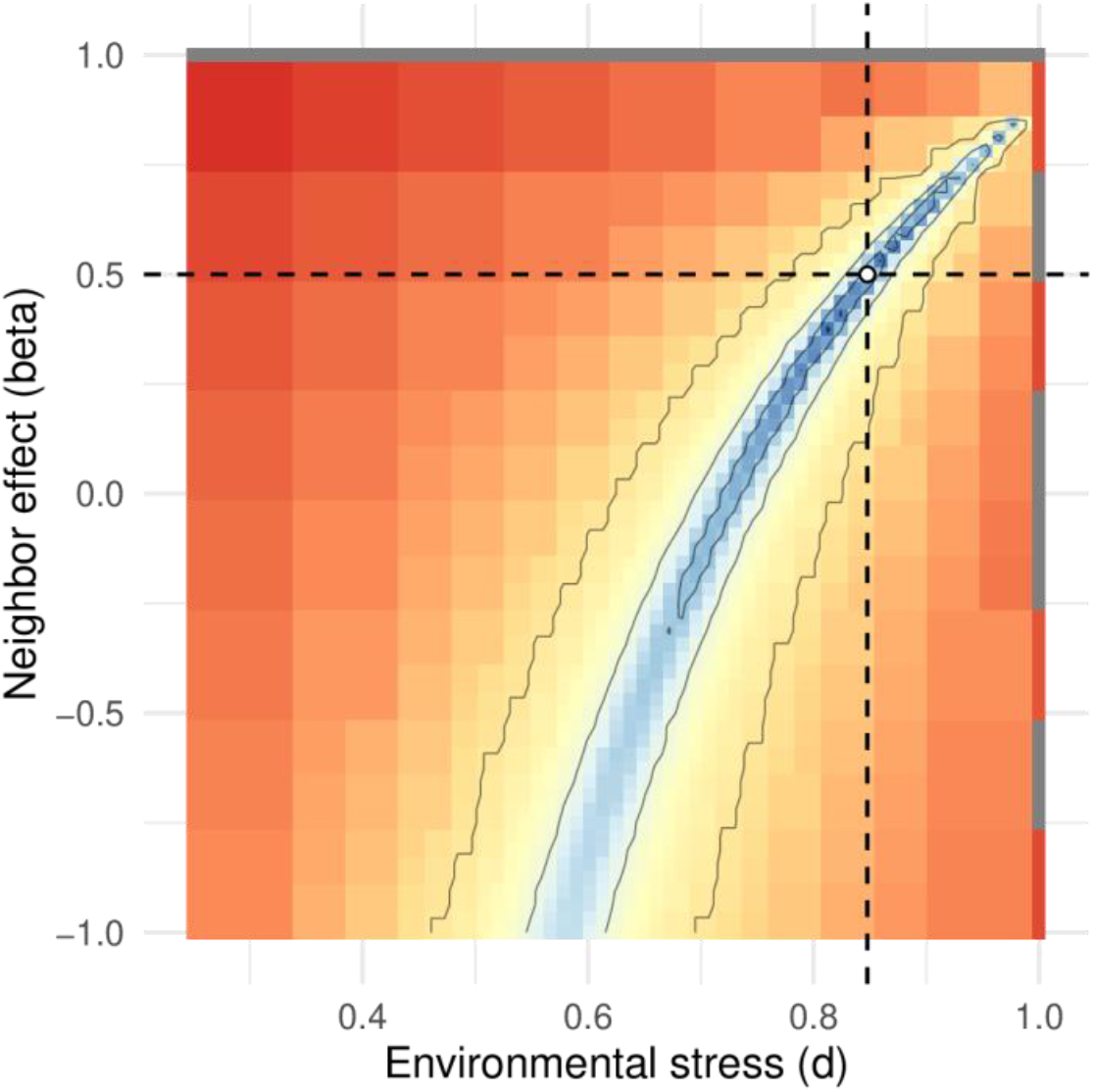
Likelihood surface as a function of the two model parameters d and β. Blue values indicate parameter combinations with high likelihood. The white dot and lines indicate the estimated parameter values.

Using this type of simulation-based inference is very expensive computationally, as this simple test requires running around a thousand simulations. This would require long-running computations with usual SCA implementations, but takes just under five minutes with `chouca` on a 2018 laptop computer (8 cores). This way of calibrating models has seldom been used in spatial ecology – a fast SCA implementation is essential to make it more accessible and test its relevance to real-world datasets.

## Conclusion

choucais an easy-to-use package to model, simulate, and visualize SCA in a reproducible way, that enables an interactive design and revision of models as well as novel methodological approaches. `chouca` does not support all types of cellular automata, and does not replace more generic modelling frameworks such as `NetLogo` (Wilensky, 1999) or `simecol` (Petzoldt & Rinke, 2007), but allows very efficient simulation for the types of SCA it supports. The package focuses on processes defined at the level of a cell, while a large set of SCA define processes at the level of pair of cells, a model representation other software packages can provide (*e*.*g*. `CellLab-CTS;` Tucker et al., 2016). Another limitation is the use of rectangular grids, which may produce discretisation artefacts that are not present in real-world patterns – future improvements may include the ability to run simulations on triangular or hexagonal grids. It is important to note that because `chouca` splits the definition of an SCA model from its simulation phase, it may use different backends for simulation. This opens the possibility of future improvements to already-implemented models without modification of existing code.

Because of the relatively low bar of entry of SCA compared to other forms of spatial modelling, we hope this work will contribute to make more accessible the testing of hypotheses linking spatial processes to patterns in ecology, as well exploring system-level consequences of specific local management decisions. `chouca` is available and maintained on CRAN, and welcomes comments, feedback and bug reports on its home page at https://github.com/alexgenin/chouca.

## Supporting information

Supplementary material 1: code for visualizing epidemic model

Supplementary material 2: code for likelihood-based inference

## Data, script and code availability

All code used for this work is freely-available at Zenodo under a CC-BY license (Génin, 2024), along with `chouca` version v0.1.99 used at the time of writing.

## Acknowledgements

This manuscript has received very constructive comments and feedback from Samuel Alizon, Broder Breckling and one anonymous reviewer, to whom we are thankful.

