## Supplementary material 1: code for visualizing epidemic model for "Easy, fast and reproducible Stochastic Cellular Automata with chouca": visual_exploration.R

```
#
# This script comes with the publication "Fast, reproducible and easy Stochastic
#   Cellular Automata with chouca", A. Génin et al. 2023
#

#
# This example illustrates how to visualize the results of an SCA using
#   chouca. We use the epidemiologic model described in Keeling and Matthew,
#   2000 (see complete reference below). We run the model on a grid, and
#   use built-in chouca functions to display the changing landscape as the
#   simulation is being run.
#
# Keeling, Matthew J. 2000. “Evolutionary Dynamics in Spatial Host–Parasite Systems.” In
# The Geometry of Ecological Interactions, edited by Ulf Dieckmann, Richard Law, and Johan
# A. J. Metz, 1st ed., 271–91. Cambridge University Press.
#
#
#

# Load the package
library(chouca)

# Set the random seed for reproducibility
set.seed(123)

# Model definition
#
# See p273 of the following book chapter:
# Keeling, Matthew J. 2000. “Evolutionary Dynamics in Spatial Host–Parasite Systems.” In
# The Geometry of Ecological Interactions, edited by Ulf Dieckmann, Richard Law, and Johan
# A. J. Metz, 1st ed., 271–91. Cambridge University Press.
# https://doi.org/10.1017/CBO9780511525537.018.
mod <- camodel(
  transition(from = "empty", to = "host",
             ~ 1 - ( 1 - g )^(4*q["host"]) ),
  transition(from = "host", to = "parasited",
             ~ 1 - (1 - T)^(4*q["parasited"])),
  transition(from = "parasited", to = "empty",
             ~ V),
  parms = list(g = 0.05, T = 0.55, V = 1), # model parameters
  wrap = TRUE,                             # use a toric space that wraps around
  neighbors = 4                            # use 4-way neighborhood
)

#
# We define the landscape as a 256x256 grid, fully covered by the host, but seeded with
# 10% of the cells in the parasited state.
ncols <- 256
nrows <- 256
initmm <- generate_initmat(mod, c(host = 0.9, parasited = 0.1, empty = 0),
                           nrow = ncols, ncol = nrows)

# We run the model from t = 0 to t = 1024. Here, iters defines the time steps
# for which we want output.
iters <- seq(0, 1024)

# We set simulation control options to display the landscape at every time step,
# and set the graphical parameter 'mar' to c(0, 0, 0, 0) so that the output
# plot has no margins. The 'fps_cap' here limits the frame rate, to make sure
# landscapes are not displayed too quickly and we can appreciate the patterns.
control <- list(custom_output_every = 1,
                custom_output_fun = landscape_plotter(mod,
                                                      fps_cap = 24,
                                                      mar = rep(0, 4)))

# We run the model. This will open a new graphical window
newmat <- run_camodel(mod, initmm, iters, control = control)


# We can export the result of the simulation to a video using the package
# 'animation' for example. Any other package that uses base graphics to
# capture the plot outputs would work as well.
#
# Here we tweak the simulation options so that
#    (1) we do not cap the fps of the output, we just want to write
#        plots on disk so we might as well do it as fast as possible
#    (2) we direct the output plots to the default output device, so that
#        animation::saveVideo() actually receives the data instead of
#        displaying things in an independent window.
#    (3) we disable margins so that the video has no white frame around
#        the landscape (set the graphical parameter mar to c(0, 0, 0, 0),
#        see options in help page ?par)
control_export <- control
control_export[["custom_output_fun"]] <- landscape_plotter(mod,
                                                           new_window = FALSE,
                                                           mar = rep(0, 4))
if ( requireNamespace("animation") ) {
  animation::saveVideo(
    run_camodel(mod, initmm, iters, control = control_export),
    video.name = "animated_ca.mp4",
    ani.width = ncols*2,   # video size in pixels
    ani.height = nrows*2,
    interval = 1/24     # 24 fps
  )
}
```
