## Supplementary material 2: code for likelihood-based inference for "Easy, fast and reproducible Stochastic Cellular Automata with chouca": adaptive_grid_sampling.R

```
#
#
# This function implements an adaptive grid sampling, in which high values
# of a function are sampled using a more precise grid than lower values. This is 
# typically useful when finding minima of likelihood surfaces.
# 
# Note that this function has been whipped up quickly and has not received
#   extensive testing outside of the code shipped with this file.
# 
# Arguments are:
#   - lower: the lower bound of parameters
#   - upper: the upper bound of parameters. lower and upper must be numerical
#       vectors with same length
#   - fun: the function to sample
#   - start_npts: the number of samples to take per parameter for the initial
#       grid
#   - prop_split: the proportion of values to split into smaller grids. At
#       each iteration of the splitting, the higher prop_split % of values will
#       be selected and splitted into subgrids
#   - max_split: the number of iterations to use when splitting. You probably
#       want to keep this low.

# Example with the banana function
#
# banana <- function(xy, a = 1, b = 100) {
#   x <- xy[1]
#   y <- xy[2]
#   # https://en.wikipedia.org/wiki/Rosenbrock_function
#   ( a - x )^2 + b * (y - x^2)^2
# }
#
# samples <- adaptive_grid_sampling(lower = c(-2, -1),
#                                   upper = c(2, 3),
#                                   fun = banana,
#                                   start_npts = 33,
#                                   prop_split = 0.10,
#                                   max_split = 4)
#
# library(ggplot2)
# ggplot(samples, aes(x = Var1, y = Var2)) +
#   geom_raster(aes(fill = ecdf(value)(value))) +
#   geom_contour(aes(z = log10(value)),
#                alpha = .4, 
#                color = "black") +
#   scale_fill_distiller(palette = "Spectral") + 
#   labs(x = "x", y = "y")

adaptive_grid_sampling <- function(lower,
                                   upper,
                                   fun,
                                   start_npts = 9,
                                   prop_split = 0.50,
                                   max_split = 3) {
  require(future.apply)
  # Make sure start_npts is odd
  start_npts <- ifelse(start_npts %% 2 == 0, start_npts + 1, start_npts)

  # assume lower/upper have same length

  # Get cell sizes
  npars <- length(upper)

  # Sample initial grid
  grid <- lapply(seq_along(lower), function(i) {
    seq(lower[i], upper[i], length.out = start_npts)
  })
  grid <- as.matrix(do.call(expand.grid, grid))
  pieces <- seq.int(nrow(grid))

  total_eval <- nrow(grid)

  message("Computing initial grid...")
  vals <- future_lapply(pieces, function(i) {
    fun(grid[i, ])
  }, future.seed = TRUE)
  vals <- unlist(vals)

  cell_sizes <- matrix(rep((upper - lower)/(start_npts-1), nrow(grid)),
                       nrow = nrow(grid), ncol = ncol(grid), byrow = TRUE)
  to_recompute <- rep(TRUE, length(vals))

  for ( split in seq.int(max_split) ) {
    message(sprintf("Starting pass %s...", split))

    # Get cells to split and split grid. We select the prop_split % of the
    # values that were just recomputed
    to_split <- is.finite(vals) & vals < quantile(vals[to_recompute], 
                                                  prop_split, na.rm = TRUE)

    # Split things
    splits <- lapply(seq_along(to_split), function(i) {

      new_lower <- grid[i, ]
      new_upper <- grid[i, ] + cell_sizes[i, ] / 2
      subgrid <- lapply(seq_along(new_lower), function(i) {
        seq(new_lower[i], new_upper[i], length.out = 2)
      })
      subgrid <- as.matrix(do.call(expand.grid, subgrid))


      # Split subgrid size
      cell_size <- matrix(new_upper - new_lower,
                          nrow = nrow(subgrid), ncol = ncol(subgrid),
                          byrow = TRUE)

      # Split vals
      if ( to_split[i] ) {
        subgrid_vals <- rep(NA, nrow(subgrid))
        subgrid_vals[1] <- vals[i] # this is the original point
      } else {
        subgrid_vals <- rep(vals[i], nrow(subgrid))
      }

      # Handle stuff outside bounds
      is_out <- rep(FALSE, nrow(subgrid))
      for ( i in seq.int(ncol(subgrid)) ) {
        is_out <- is_out | ( subgrid[ ,i] < lower[i] | subgrid[ ,i] > upper[i] )
      }

      list(subgrid[!is_out, ], cell_size[!is_out, ], subgrid_vals[!is_out])
    })

    grid <- do.call(rbind, lapply(splits, function(o) o[[1]]))
    cell_sizes <- do.call(rbind, lapply(splits, function(o) o[[2]]))
    vals <- do.call(c, lapply(splits, function(o) o[[3]]))

    # Recompute values
    to_recompute <- is.na(vals)
    total_eval <- total_eval + sum(to_recompute)
    message(sprintf("Recomputing %s values...", sum(to_recompute)))
    new_vals <- future_lapply(which(to_recompute), function(i) {
      fun(grid[i, ])
    }, future.seed = TRUE)
    vals[to_recompute] <- unlist(new_vals)

  }

  message(sprintf("Done, %s function evaluations used", total_eval))
  # Return the grid and values
  data.frame(grid, value = vals)
}
```
