## Supplementary material 2: code for likelihood-based inference for "Easy, fast and reproducible Stochastic Cellular Automata with chouca": local_inter_inference.R

```
#
# This script comes with the publication "Fast, reproducible and easy
#   Stochastic Cellular Automata with chouca", A. Génin et al. 2023
#
#
# This example illustrates the inference of interactions between
#   habitat-forming organisms on simulated data, with the example of inferring
#   plant-plant interactions from a landscape snapshot.
#
#

# For now we need to install the development version of this package. We install 
# a specific commmit for better reproducibility
devtools::install_github("spatial-ews/spatialwarnings", 
                         ref = "ea055d0688c888e323def11fbddf4cb001b8bf13")

# We load the required packages for this session
library(plyr)
library(spatialwarnings)
library(ggplot2)
library(future.apply)
library(chouca)

# Load helper function
source("./adaptive_grid_sampling.R")

# Set the random seed for reproducibility
set.seed(123)

# Set the size of the grid the SCA is going to run on
ncols <- 256
nrows <- 256

# Here we define the SCA model of bare/plant. With two states (bare/plant)
# and two transitions (bare to plant and vice versa). This model has two
# parameters, that are provided with default values using the 'parms'
# argument to camodel. We use a toric space (i.e. leftmost/topmost cells are
# neighbors of the right/bottom row of the lattice), and a 4 neighbor geometry
# (von-Neumann neighborhood).
mod_on_recruitment <- camodel(
  transition(from = "bare", to = "plant",
             ~ p["plant"]),
  transition(from = "plant", to = "bare",
             ~ d * ( 1 - beta * q["plant"] ) ),
  parms = list(beta = 0.1,
               d = 0.05),
  wrap = TRUE,
  neighbors = 4
)

# This example simulates observed data, then tries to recover the parameters used to
# simulate the data. Here, we define the default model parameters beta and d used to
# simulate the observed data. These are the values we are going to try to
# recover using the likelihood-based approach.
#
beta_simu <- .50
d_simu <- 0.85

# We create a large initial matrix of edge size 4*nrows*ncols, randomly
# filled with 50% plants and 50% bare cover. This mimicks what would be
# obtained from a remote sensing image.
initmm <- generate_initmat(mod_on_recruitment,
                           c(plant = 0.5, bare = 0.5),
                           nrows*4, ncols*4)


# SIMULATE OBSERVED DATA
# ----------------------

# To simulate observed data, we will run the model for 1024 iterations, which is more
# than enough to reach equilibrium here
iters <- seq(0, 1024)

# This is the 'control' list to run the simulation.
control <- list(
  # We save covers (% of the lanscape in each state) at each iteration
  save_covers_every = 1,
  # We save the full landscape (a snapshot) only on the last iteration
  save_snapshots_every = max(iters),
  # We use the 'compiled' engine to generate C++ code and compile it on the
  # fly. This is much faster than using the default "cpp" backend
  engine = "compiled",
  # We use only one core. Because this simulation is run on a 
  # relatively small grid and using memoization, parallelizing 
  # computations over multiple cores is counter-productive
  # here. We will parallelize at a higher level.
  cores = 1,
  # We decompose time steps into substeps so that probabilities are never
  # close to one
  substeps = 2,
  # Because this is a model with a low number of state (2), it is worth
  # precomputing probabilities for all neighbor configurations instead of
  # recomputing them for all cells.
  precompute_probas = TRUE
)

# We update the model definition with the parameter values used to generate the
# simulated empirical data.
mod_on_recruitment <- update(mod_on_recruitment,
                             parms = list(d = d_simu,
                                          beta = beta_simu))

# Run the model
newmat <- run_camodel(mod_on_recruitment, initmm, iters, control = control)

# Extract the final landscape of the simulation, then compute the observed distribution
# of pairs of cells. This uses the function pair_counts() from package spatialwarnings.
# Here this distribution is the count of each pairs of possible pairs found in the
# landscape, which is a set of three integers for the pairs (plant-plant, plant-empty and
# empty-empty).
obs_landscape <- tail(newmat[["output"]][["snapshots"]], 1)[[1]]

pcounts_obs_on_recruitment <- pair_counts(obs_landscape,
                                          wrap = FALSE,
                                          use_8_nb = FALSE,
                                          prop = FALSE)


# This function will be used to run the model when finding parameters that are
# closest to empirical patterns. Argument 'theta' is a vector of two values
# containing first the mortality rate (d), and second the neighbor effect
# (beta). The 'mod' argument here is used to pass the 'mod_on_recruitment' object, which
# is the 'camodel' object defined above (this makes parallelizing easier).
#
# This function runs the model for a number of iterations to allow reaching
# equilibrium, and saves the distribution of pairs of cells for this simulation.
#
# It also computes the likelihood given the parameter set 'theta' and the observed
# distribution of cell pairs (defined above in pcounts_obs_on_recruitment). We assume
# that the distribution of pairs follows a multinomial distribution of
# parameters:
#    mu = (mu_pp, mu_p0, mu_00) which are the probabilities of drawing a pair of a
#      given type (plant-plant or pp, plant-empty or p0, empty-empty or 00), and
#    N = the total number of pairs ('tries' in binomial terms) in the landscape (this is
#      a fixed number and only depends on the size of the landscape, the neighborhood
#      type and whether we wrap around the edges or not
#
# It is important to note that we use a stochastic simulation to estimate the likelihood,
# so there may be some variation in its value, even for the same set of parameters theta. 
# Here, we estimate the likelihood using the simulated landscape at each 
# iteration of the model after a burn-in period. We run the model for 512 
# iterations and use 256 iterations for burn-in, so each call to run_simu() returns 
# 256 estimates (512 minus 256) of the likelihood. It is summarized into a single 
# value by the function get_ll() below. 
# 
run_simu <- function(theta, mod) {

  # We extract the parameters from the theta vector, and update the model definition
  # with the new parameter values.
  d <- theta[1]
  beta <- theta[2]
  mod <- update(mod, list(d = d,
                          beta = beta),
  # Here we signify that we want to check the model using all possible neighbor
  # configurations. Warnings will be produced, e.g. if probabilities can reach
  # value above one, or below zero. Use "quick" or "none" to speed this up.
                check_model = "quick")

  # Define the vector of iterations for which we want model output
  iters <- seq(0, 2^9) # 512 time steps

  # Disable output
  control[["console_output_every"]] <- 0
  control[["save_snapshots_every"]] <- max(iters)
  control[["save_covers_every"]] <- 0

  # Generate an initial landscape, then run the model once parameters and functions
  # are set up
  initmm <- generate_initmat(mod, c(0.5, 0.5), nrows/2, ncols/2)
  
  # We define a function to count pairs of cells as the simulation is running, but 
  # not before a burn-in period
  compute_ll_fun <- function(t, mat) { 
    
    # If we are before the end of burn-in period, do nothing and return early
    if ( t < max(iters)/2 ) { 
      return(NULL) 
    }
    
    # Compute pair distribution
    pairs <- pair_counts(mat,
                         wrap = FALSE,
                         use_8_nb = FALSE,
                         prop = FALSE)
    
    # Format observed and current pair distribution
    obs <- pcounts_obs_on_recruitment
    obs <- obs[lower.tri(obs, diag = TRUE)]
    pc <- pairs
    pc <- pc[lower.tri(pc, diag = TRUE)]
    
    # Compute the likelihood
    ll <- dmultinom(obs, size = sum(obs), prob = pc / sum(pc), log = TRUE)
    
    return(ll)
  }
  
  control[["custom_output_fun"]] <- compute_ll_fun
  control[["custom_output_every"]] <- 1
  
  run <- run_camodel(mod, initmm, iters, control = control)
  
  # Summarize distribution of pairs
  # This can produce non-integer numbers of pairs but that's ok for what we want to 
  # do next
  lls <- Filter(function(o) ! is.null(o), run[["output"]][["custom"]])
  
  # Return the computed quantities (here the likelihoods)
  return(
    list(lls = lls)
  )
}

# run_simu(c(d_simu, beta_simu), mod_on_recruitment)


# run_simu() returns a set of estimates of the likelihood, that we need to further 
# summarize into a single value. We return the most frequent value of the likelihood 
# out of those returned by run_simu(), estimated as the maximum of a kernel-estimated 
# density. 
get_ll <- function(theta, mod, ...) { 
  
  # Run the simulation, and extract estimates of the likelihood
  run <- run_simu(theta, mod)
  lls <- unlist(run[["lls"]])
  
  # Sometimes log likelihood is infinite, and density() will error out -> handle 
  # this case. 
  if ( ! sum(is.finite(lls)) > 2 ) {
    return(-Inf)
  }
  
  # Use most frequent ll value
  d <- density(lls)
  with(d, x[y == max(y)])
}

# get_ll(c(0.5, 0.5), mod = mod_on_recruitment)

# Sample parameters to build the likelihood surface
plan(multisession) # create workers for parallel processing
system.time({
  res <- adaptive_grid_sampling(lower = c(0.25, -1), # upper and lower bounds
                                upper = c(1, 1),     # on parameters
                                function(theta) {
    # Return negll for minimization purposes
    - get_ll(theta, mod_on_recruitment)
  }, start_npts = 8, prop_split = 0.50, max_split = 3)
})
plan(sequential) # destroy workers
# 239.658 seconds on t480 ~= 4 minutes

values_format <- res
names(values_format) <- c("d", "beta", "negll")
best <- subset(values_format, negll == min(negll) & is.finite(negll))

# Display best estimate and LL surface
grid_plot <- ggplot(values_format,
                    aes(x = round(d, 8),
                        y = round(beta, 8))) +
  # Show likelihood surface
  geom_raster(aes(fill = negll)) +
  geom_contour(aes(z = negll - best[ ,"negll"]),
               breaks = c(-Inf, 1, 10, 100, 1000, 10000, 100000, Inf),
               color = "black", linewidth = 0.2, alpha = .5) +
  # Show best combination
  geom_hline(aes(yintercept = beta), data = best, 
             linetype = "dashed") + 
  geom_vline(aes(xintercept = d), linetype = "dashed",
             data = best) +
  geom_point(aes(x = d, y = beta),
             data = best, 
             pch = 21, fill = "white") +
  scale_fill_distiller(palette = "RdYlBu",
                       trans = "log10") +
  theme_minimal() +
  theme(legend.position = "none") +
  labs(x = "Environmental stress (d)",
       y = "Neighbor effect (beta)")

grid_plot

ggsave("./inter_inference.pdf", width = 4, height = 4)
```
