## Supplementary figures and images for "Easy, fast and reproducible Stochastic Cellular Automata with chouca"

### inter_inference.pdf

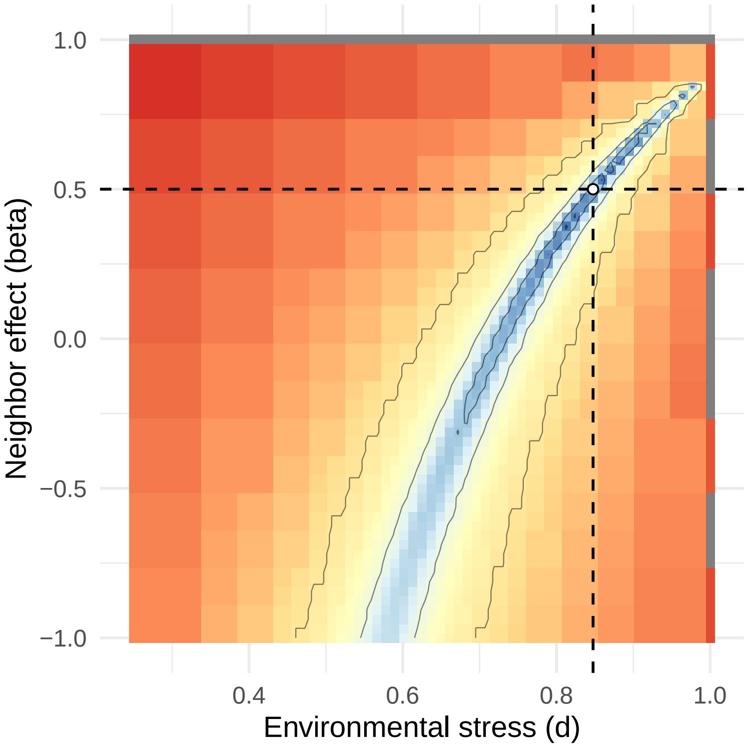

### inter_inference.png

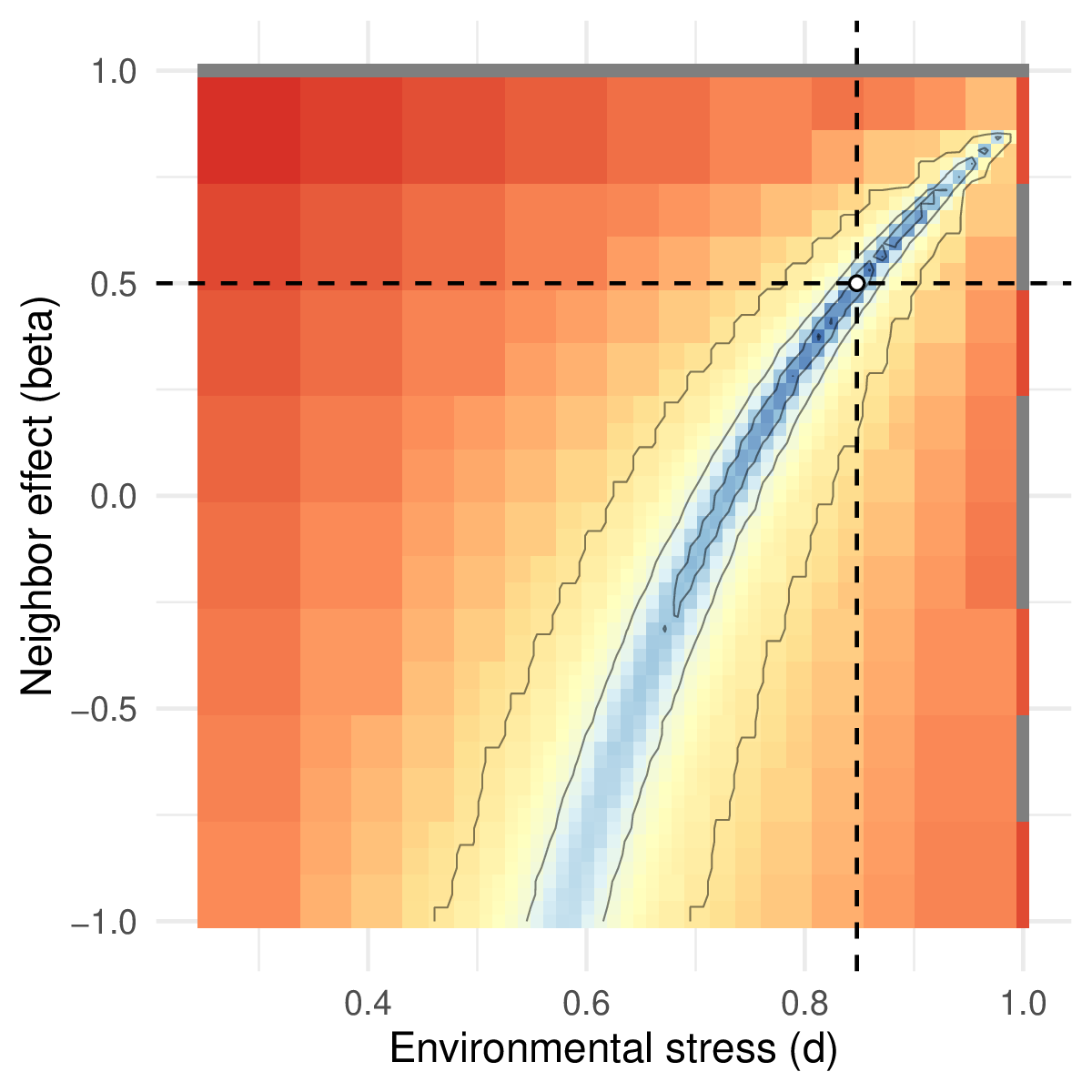
